# Investigating the potential role of capsaicin in facilitating the spread of coxsackievirus B3 via extracellular vesicles

**DOI:** 10.1101/2024.12.02.626352

**Authors:** Shruti Chatterjee, Haylee Tilley, Devin Briordy, Richard T. Waldron, Ramina Kordbacheh, William D. Cutts, Alexis Cook, Stephen J. Pandol, Brandon J. Kim, DeLisa Fairweather, Jon Sin

## Abstract

Coxsackievirus B3 (CVB3) is a non-enveloped picornavirus that can cause systemic inflammatory diseases including myocarditis, pericarditis, pancreatitis, and meningoencephalitis. We have previously reported that following infection, CVB3 localizes to mitochondria, inducing mitochondrial fission and mitophagy, while inhibiting lysosomal degradation by blocking autophagosome-lysosome fusion. This results in the release of virus-laden mitophagosomes from the host cell as infectious extracellular vesicles (EVs) which allow non-lytic viral egress.

Transient receptor potential vanilloid 1 (TRPV1/*TRPV1*) is a heat and capsaicin-sensitive cation channel that regulates mitochondrial dynamics by inducing mitochondrial membrane depolarization and fission. In this study, we found that treating cells with the TRPV1 agonist capsaicin dramatically enhances CVB3 egress via EVs. Analysis of the released EVs revealed increased levels of viral capsid protein VP1/*VP1*, mitochondrial protein TOM70/*TOMM70*, and fission protein phospho-DRP1/*DNM1L* (Ser 616). Moreover, these EVs exhibited increased levels of heat shock protein HSP70/*HSPA1A*, suggesting a potential role of these chaperones in facilitating infectious EV release from cells.

Furthermore, TRPV1 inhibition with capsazepine significantly reduced viral infection *in vitro*. We previously observed similar effects *in vitro* with another TRPV1 inhibitor SB-366791. Our current *in vivo* studies found that SB-366791 significantly mitigates pancreatic damage and reduces viral titers in mouse model of CVB3 pancreatitis.

Given the lack of understanding regarding the factors that contribute to diverse clinical manifestations of CVB3, our study highlights capsaicin and TRPV1 as potential exacerbating factors that facilitates CVB3 dissemination via mitophagy-derived EVs.

**IMPORTANCE:** CVB3 is a prevalent pathogen responsible for a range of severe diseases, including myocarditis, pericarditis, pancreatitis, and meningoencephalitis. Despite its clinical significance, factors that determine the severity of CVB3 infection and why some individuals experience life-threatening manifestations while others have mild, cold-like symptoms remain poorly understood. This study provides new insights into the molecular mechanisms underlying CVB3 dissemination and pathogenesis. By investigating the role of capsaicin, a common dietary component, in modulating viral spread, we demonstrate that activation of TRPV1 by capsaicin enhances release of infectious CVB3 via mitophagy-derived EVs. Our results offer novel evidence that modulating TRPV1 activity could influence the clinical outcomes of CVB3 infection, opening new avenues for therapeutic interventions. Given the widespread consumption of capsaicin, this study highlights an important dietary factor that could play a role in shaping CVB3 pathogenesis and its clinical manifestations, underscoring the potential for targeted strategies to mitigate severe disease outcomes.

## INTRODUCTION

Coxsackievirus is a positive-sense single stranded RNA virus that belongs to the *Enterovirus* genus within the *Picornaviridae* family [1]. These non-enveloped viruses can infect a range of tissues, notably the central nervous system, heart, and pancreas, causing diseases from mild febrile illnesses to severe conditions like myocarditis, pericarditis, meningitis, and pancreatitis [2, 3]. Their impact is particularly pronounced in neonates and young children; however, adults are also susceptible to these diseases [4, 5]. Coxsackieviruses can be categorized into two primary groups: coxsackievirus A (CVA) and coxsackievirus B (CVB). CVA primarily targets the skin and mucous membranes, often causing conditions like hand, foot, and mouth disease (HFMD) and herpangina, while CVB is known to target internal organs, including the heart, pancreas, and central nervous system [6–8]. CVB can further be subdivided into six distinct serotypes, labeled CVB1 through CVB6, with CVB3 being particularly significant due to its impact on human health and its role in a range of serious diseases [9–13]. Currently, whether CVB3-infected individuals will suffer lethal manifestations as opposed to a mild, cold-like symptomatology remains unclear. Further research into mechanisms that enhance viral infection could shed light on environmental contributors to differential clinical presentations.

Our previously published studies suggest that CVB3 can subvert mitophagy in host cells– a cellular degradation pathway by which damaged or dysfunctional mitochondria are packaged within autophagosomes that eventually become degraded via their fusion with acidic lysosomes [14–17]. A key process required for mitophagy is mitochondrial fission which removes damaged mitochondria from the healthy mitochondrial network [18]. In our earlier studies, we showed that CVB3 virions not only induce mitophagy, but also becomes incorporated into mitophagosomes. Furthermore, others have shown that CVB3 can impair autophagosome-lysosome fusion, thus permitting virus-laden mitophagosomes to be released from the host cell as infectious extracellular vesicles (EVs) [19–23]. This mechanism serves as a non-lytic mode of viral dissemination that prolongs replication within host cells and enables them to evade host neutralizing antibodies. We have previously investigated transient receptor potential (TRP) channels to understand their potential roles to mediate CVB3-induced mitophagy and mitochondrial fission associated with CVB3 infection. Our previous studies have explored the relationship between TRP channels and CVB3 infection, specifically in regard to transient receptor potential melastatin 8 (TRPM8). We have reported that the cold and menthol-sensitive TRPM8 plays a crucial role in inhibiting mitochondrial fission (a prerequisite for mitophagy) during CVB3 infection [22]. Moreover, activating TRPM8 with the cooling compound menthol produced significantly antiviral effects. To further explore the involvement of TRP channels in CVB3 infection, in this current study we investigated another TRP channel, TRPV1, to determine its potential significance in CVB3 infection. TRPV1 is a heat and capsaicin-activated ion channel that has previously been shown to induce mitochondrial membrane depolarization and subsequent mitochondrial fission following its activation; the opposite of what has been observed in TRPM8 [24–26]. The connection between CVB3-induced proviral mitochondrial fission and TRPV1-induced fission suggests that TRPV1 could play an intermediary role in increasing CVB3 infection/dissemination.

In this study, we observed that activating TRPV1 with the commonly consumed spicy compound capsaicin significantly increased CVB3 infection in HeLa cervical cancer cells. This was mediated by dramatically increased viral shedding during capsaicin exposure. When interrogating the composition of released EVs we not only saw significantly more viral capsid protein VP1, but also observed increased levels of mitochondrial protein TOM70 and mitochondrial fission protein phospho-DRP1 (Ser 616). These findings suggest the potential role of capsaicin in inducing proviral mitophagy which facilitates EV-based viral spread. In addition, capsaicin treatment further enhanced the expression of heat shock protein HSP70 in EVs. Numerous studies have demonstrated that HSP70 is a proviral protein in the context of CVB3 infection as it facilitates viral protein translation both at the initiation and elongation stages [27]. The heat shock response in cells is initiated by an increase in the fluidity of specific membrane domains and earlier studies have shown that plasma membrane-associated TRPV1 can directly trigger the membrane-dependent activation of HSPs [28, 29]. Given that HSP70 has been identified as a potential biomarker of EVs shed from certain diseased cell types such as cancer cells, we hypothesize that capsaicin not only promotes mitophagy-based viral EV release, but more specifically facilitates the biogenesis of HSP70-positive viral EVs. This was further corroborated by silencing *HSPA1A* in HeLa cells, which as associated with (or led to) a drastic reduction in viral protein concentration within EVs. All these findings reveal how capsaicin-induced TRPV1 activation not only amplifies CVB3 infection but also highlights a novel mechanism of viral spread through HSP70-enriched EVs, unveiling new targets for therapeutic intervention. Furthermore, the proviral action of TRPV1 was validated *in vitro* using the competitive antagonist capsazepine and *in vivo* using the allosteric inhibitor SB-366791 which not only demonstrated reduced viral infection, but also mitigated pancreatic viral burden, inflammation and damage.

Knowing that capsaicin exacerbates CVB3 infection as well as viral release via EVs, we hypothesize that this common dietary component may predispose to severe CVB3 infections. Due to the large gaps in our understanding of susceptibility factors that contribute to CVB3 diseases, we anticipate that this study will shed light on a very common dietary component that could potentially exacerbate CVB3 disease manifestation.

## RESULTS

### Capsaicin enhances CVB3 infection

Recent studies have highlighted the role of TRPV1 in airway hypersensitization and the spread of various respiratory viruses [30]. In 2020, our research established a novel link between TRPV1 and CVB3. We observed that inhibiting TRPV1 with the potent and selective antagonist SB-366791 dramatically inhibits CVB3 infection *in vitro* [22]. Based on this, we hypothesized that chemicals that activate TRPV1, such as capsaicin, might exacerbate infection by triggering mitochondrial fission. To test this hypothesis, we treated HeLa cells with 10 μM capsaicin for 15 mins before infecting them with enhanced green fluorescent protein (EGFP)-expressing-CVB3 (EGFP-CVB3) at a multiplicity of infection (MOI) of 0.001. Following infection for 48 hours, a substantial enhancement of viral EGFP expression was observed in capsaicin-treated cells using fluorescence microscopy and flow cytometry (Fig. 1 A, B and Supplemental Fig. S1). This was further confirmed by plaque assay which revealed significantly higher viral titers in the extracellular media of capsaicin-treated cells (*p* = 0.0027) (Fig. 1C). However, western blot analysis of the corresponding cell lysates revealed no significant differences in intracellular viral capsid protein VP1 levels between capsaicin-treated and vehicle-treated infected samples (*p* = 0.8227) (Fig. 1D, E). This discrepancy between western blot and plaque assay suggests that following infection for 48 hours, capsaicin may potentially induce viral egress from host cells, resulting in higher viral titers in media compared to cellular VP1 expression.

**FIG 1.**
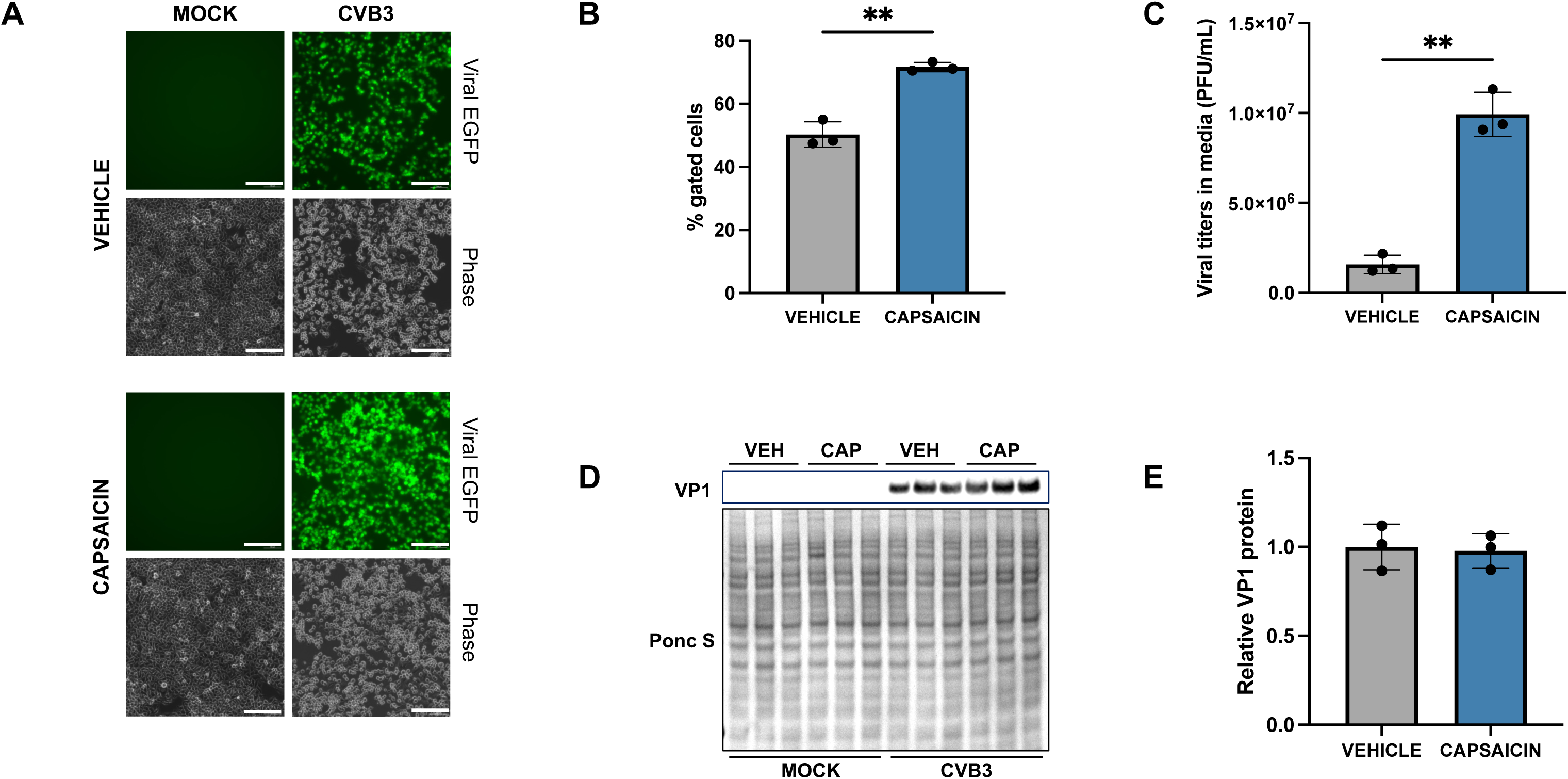
Capsaicin enhances CVB3 infection. Human cervical cancer cells HeLa were treated with 10 µM Capsaicin for 15 mins and infected with EGFP-CVB3 at multiplicity of infection (MOI) of 0.001 for 48 hours. **A**. Fluorescence microscopy compares viral EGFP expression between vehicle (DMSO-treated) and capsaicin-treated cells. Phase contrast images are shown below respective fluorescence image fields. Scale bars represent 100 µm. **B.** Flow cytometry of infected cells revealed significantly increased EGFP-positivity in capsaicin-treated cells as compared to controls. **C.** Plaque assay quantification of infectious virus in media from cells in A. **D.** Western blot on cell lysates from A showing VP1 viral capsid protein. Ponceau S is shown below. (VEH: Vehicle; CAP: Capsaicin). **E.** Densitometric quantification of western blot in B (** p< 0.01; Student’s t-test, n = 3, error bars represent standard deviation).

### Capsaicin induces viral egress and EV-mediated spread of CVB3

In recent years, several groups including ourselves have demonstrated that non-enveloped viruses like CVB can spread from one cell to another via extracellular vesicles or EVs [15, 23, 30, 31]. This strategy of cellular escape has been proposed to confer a number of advantages including prolonging viral replication, transfer of numerous virions in each EV, and evasion from host adaptive immunity [15, 21, 23]. To investigate the impact of capsaicin on the egress and dissemination of CVB3 via EVs. HeLa cells were treated with capsaicin as previously described, and EVs were isolated from the extracellular media after CVB3 infection for 48 hours. Western blot analysis revealed that EVs from capsaicin-treated cells contained significantly higher amounts of VP1 compared to those from DMSO (vehicle)-treated cells (*p* = 0.0385) (Fig. 2A, B). Notably, VP1 levels within EVs were significantly greater than that observed in cell lysates (*p* = 0.8227), as shown in Figure 1D. Next, we showed that the infectious viral titers were significantly higher in EVs isolated from capsaicin-treated cells compared to those from the vehicle-treated cells (*p* = 0.0015) (Fig. 2C). In contrast, we found that the change in the amount of free virus present in the supernatant of capsaicin-treated cells was not significantly different from the vehicle-treated cells (*p* = 0.8230). Taken together, these results strongly indicate that capsaicin treatment promotes the release of infectious EVs from host cells.

**FIG 2.**
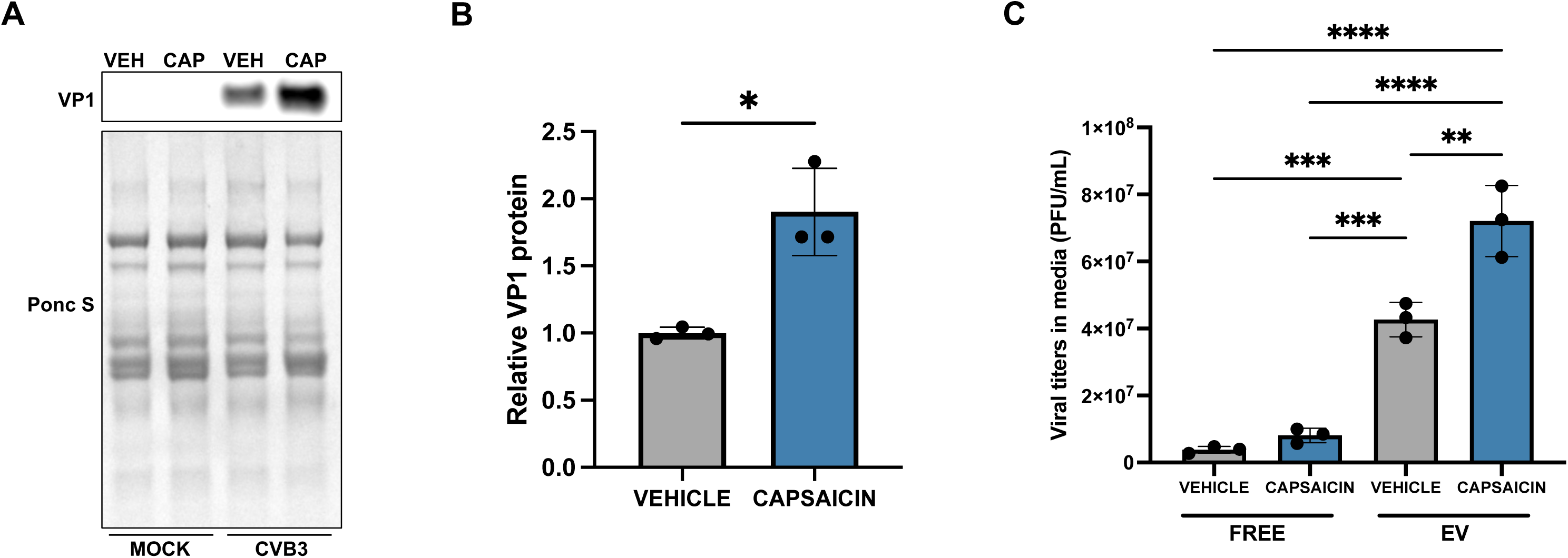
Capsaicin induces viral release in EVs. HeLa cells were treated with 10 µM Capsaicin for 15 mins and infected with EGFP-CVB3 at an MOI of 0.001 for 48 hours. **A.** Western blot on isolated EVs showing VP1 viral capsid protein. Ponceau S is shown below. (VEH: Vehicle; CAP: Capsaicin) **B.** Densitometric quantification of western blot in A. **C.** Plaque assay of EV versus free virus in capsaicin and DMSO-treated cells (* p< 0.05, ** p< 0.01; *** p<0.001; **** p<0.0001; Student’s t-test, n = 3).

### Capsaicin increases mitochondrial content in EVs

To further investigate the origin of these EVs, we analyzed them for mitochondrial markers. HeLa cells were pretreated with 10 μM capsaicin for 15 minutes before infection with EGFP-CVB3 at an MOI of 0.001. After 48 hours, we isolated EVs and performed western blot analysis to examine the mitochondrial protein TOM70 and mitochondrial fission protein phospho-DRP1 (Ser 616) (Fig. 3 A-C). As expected, CVB3 infection promoted a dramatic increase in the levels of these proteins in EVs. However, capsaicin treatment by itself also led to a significant increase in phospho-DRP1 (Ser 616) (*p* = <0.0001) and TOM70 (*p* = 0.0052) levels within EVs, implying that the TRPV1 agonist promotes mitochondrial fission and contributes to the presence of mitochondrial fragments within EVs. Furthermore, combining capsaicin treatment with infection enhanced the expression of these proteins even more, albeit modestly (*p* = <0.0001 for phosphor-DRP1 and 0.0016 for TOM70). Since intracellular expression of these proteins did not show significant changes, both with treatment and infection (Supplemental Fig. S2 A-C), these findings imply that capsaicin enhances CVB3 spread via mitophagosome-bound EVs.

**FIG 3.**
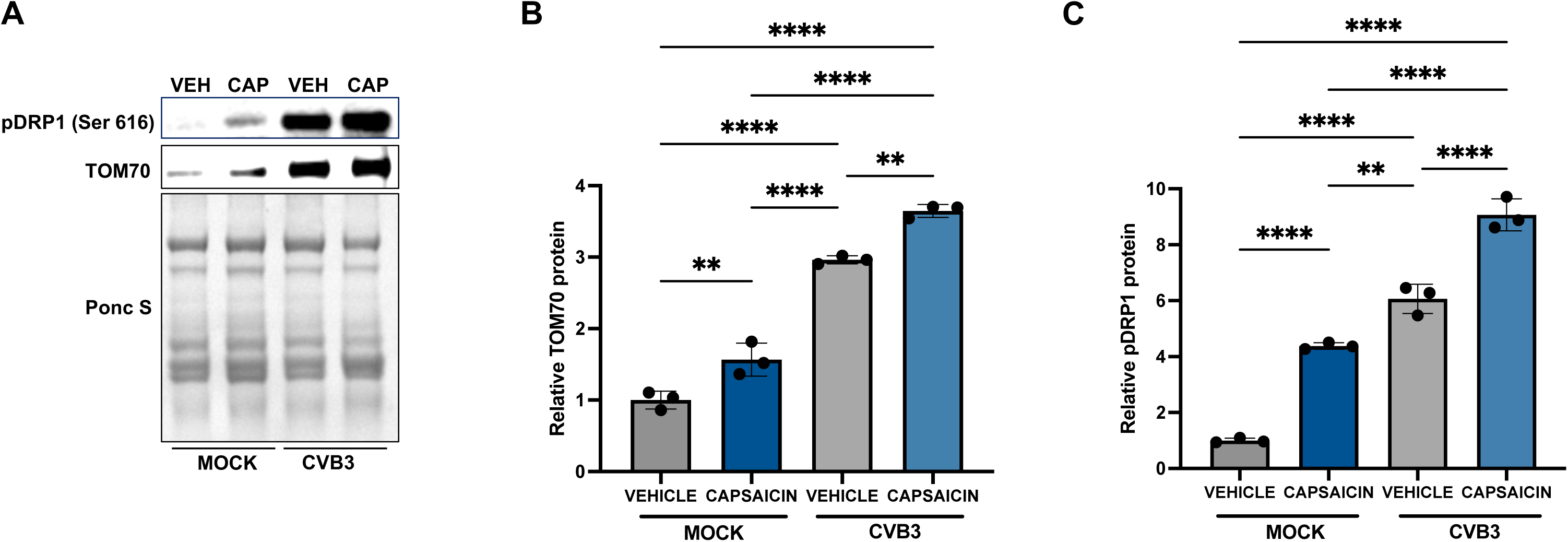
Capsaicin increases mitochondrial content in EVs. HeLa cells were treated with 10 µM Capsaicin for 15 mins and infected with EGFP-CVB3 at multiplicity of infection (MOI) of 0.001 for 48 h. **A.** Western blot on EVs detecting TOM70 and phosphor DRP-1 (Ser 616). Ponceau S is shown below. (VEH: Vehicle; CAP: Capsaicin) **B, C.** Densitometric quantification of western blots in A. (* p< 0.05, ** p< 0.01; *** p<0.001; **** p<0.0001; Student’s t-test, n = 3).

### Capsaicin-induced infectious EVs are enriched with HSP70

The 70 kDa heat shock protein (HSP70) is a key component of the cellular chaperone network, crucial for managing various cellular stresses [32] and has been considered to be a biomarker for tumor-derived exosomes [33–35]. Moreover, HSP70 expression can be triggered due to an increase in the fluidity of specific membrane domains in plasma membrane, and TRPV1, being a key regulator in cellular thermosensory pathway, plays a crucial role in inducing membrane-dependent HSPs [28, 29]. Although HSPs are biologically designed to maintain cellular homeostasis, viruses can exploit them for protein folding and to increase their chances of survival in adverse host conditions [36–39]. For CVB3, HSP70 has been shown to facilitate viral infection by promoting viral protein translation at both the initiation and elongation stages [27]. Moreover, HSP70 was reported to induce mitophagy by interacting with PTEN-induced kinase 1 (PINK1) in HEK 293 cells [40]. In this study, we intended to investigate the role of this chaperone in the release of CVB3-laden EVs from capsaicin-treated HeLa cells. HeLa cells were treated with 10 µM capsaicin for 15 minutes, followed by infection with EGFP-CVB3 at an MOI of 0.001 for 48 hours. Subsequently, EVs were isolated and analyzed for HSP70 presence. Western blotting revealed that viral EVs from capsaicin-treated cells contained higher levels of HSP70 compared to those treated with DMSO (Fig. 4 A, B). Although vehicle-treated infected EVs also showed increased HSP70 levels compared to mock-infected EVs, possibly due to proviral effects of HSP70 activity, capsaicin treatment in addition to infection showed a subtle but significant increase in HSP70 content in EVs (*p* = 0.0002). In contrast, changes in HSP70 expression in cell lysates were less pronounced with either capsaicin-exposure or CVB3 infection (Fig. 4 C, D). Overall, these findings suggest that capsaicin not only enhances the release of HSP70-enriched EVs, but that HSP70 may have a previously undescribed role in EV-based viral egress.

**FIG 4.**
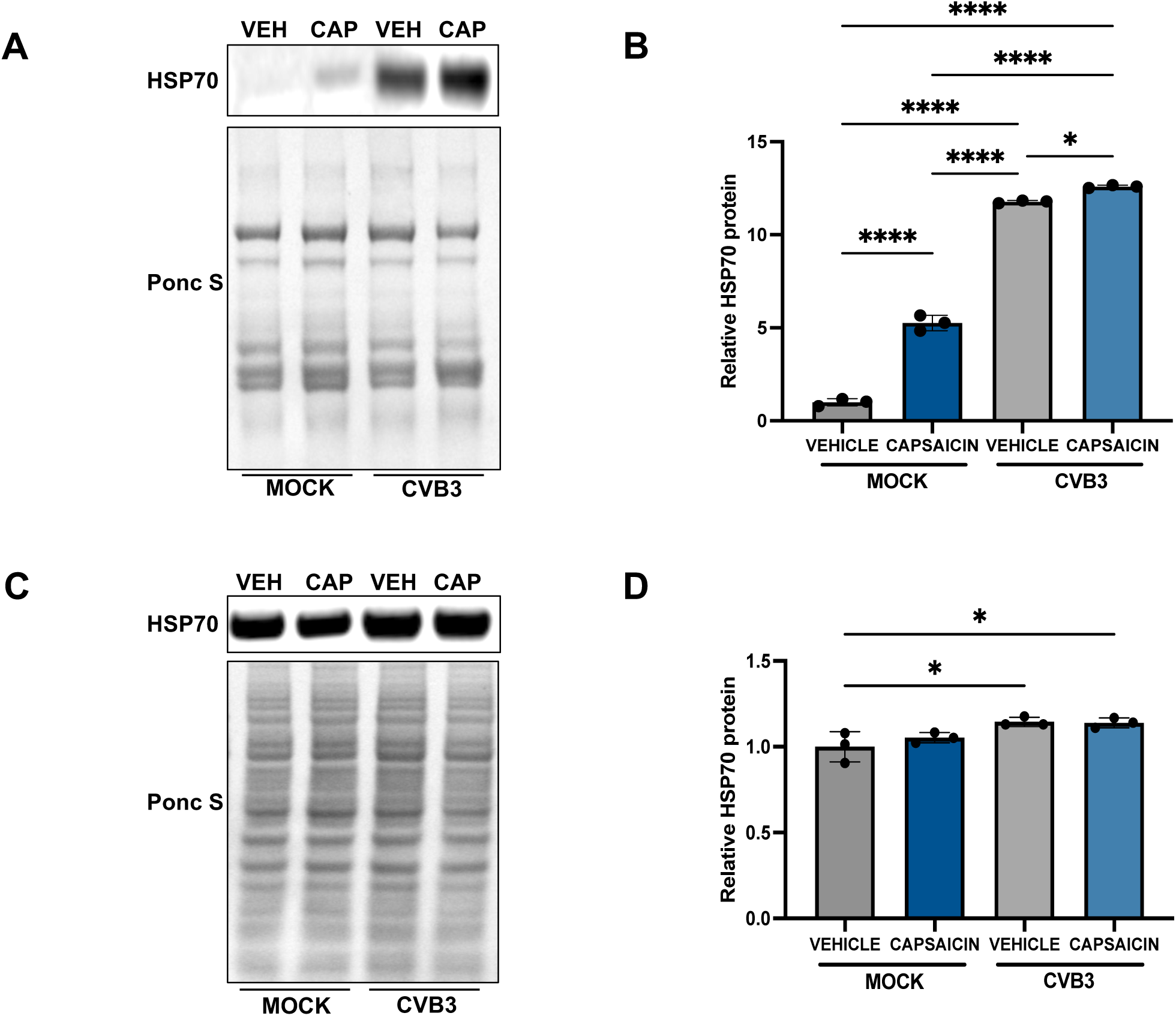
Capsaicin induced EVs are enriched with HSP70. HeLa cells were treated with 10 µM Capsaicin for 15 mins and infected with EGFP-CVB3 at MOI of 0.001 for 48 hours. **A.** Western blot on EVs and **(C)** whole cell lysates detecting HSP70. Ponceau S is shown below. (VEH: Vehicle; CAP: Capsaicin) **B, D.** Densitometric quantification of western blot in A and C. (* p< 0.05, ** p< 0.01; *** p<0.001; **** p<0.0001; Student’s t-test, n = 3).

### Silencing HSP70 inhibits CVB3 release in EVs

To further ascertain the involvement of HSP70 in dissemination of EVs from cells, we further transfected cells with an *HSPA1A*-targeting siRNA. After 48 hours of silencing, the cells were infected with EGFP-CVB3 at an MOI of 0.001. Following infection for 24 hours, EVs and cell lysates were collected, and western blot performed for both HSP70 and VP1 expression. Although the reduction in cellular HSP70 levels were somewhat modest after silencing (Fig. 5 B, C), fluorescence microscopy, western blotting of cell lysates, and plaque assays of the extracellular media demonstrated a significant reduction in viral presence in the *HSPA1A*-silenced infected cells, indicating that this level of silencing still resulted in a marked inhibition of CVB3 infection (Fig. 5 A, B, D, E). Notably, the decrease in viral titers was more pronounced than the reduction in VP1 expression in cell lysates, supporting that *HSPA1A* silencing impacts viral release and potentially reduces the release of infectious EVs. To explore this further, we analyzed VP1 expression in isolated EVs and found that *HSPA1A*-silenced EVs contained significantly lower levels of VP1 compared to control EVs (Fig. 6 A-C). These results support our hypothesis that HSP70 plays an important role in the release of EVs from infected cells, and capsaicin can enhance viral spread by increasing release of these HSP70-laden EVs. It is unclear if these EVs are derived from mitophagosomes, or from a novel parallel pathway, which like mitophagy, contributes to viral spread from one cell to another.

**FIG 5.**
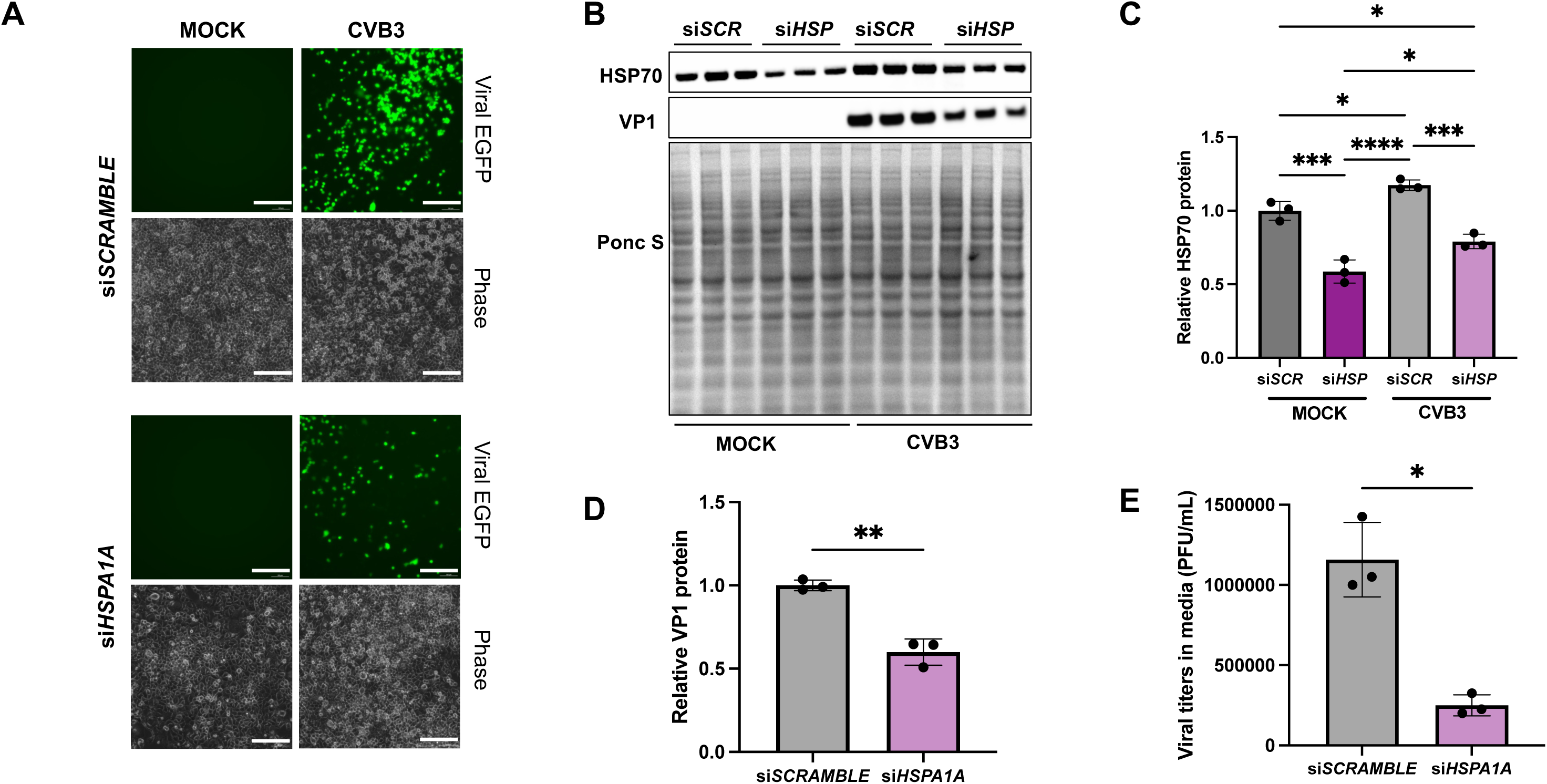
Silencing HSP70 inhibits CVB3 infection. **A.** HeLa cells were transfected with siRNA targeting HSP70 (si*HSPA1A*) or scrambled RNA (si*SCRAMBLE*) for 48 hours and subsequently infected with EGFP-CVB3 at MOI 0.001 for 24 h. **(A)** Fluorescence microscopy comparing viral EGFP expression between cells transfected with either si*SCRAMBLE* or si*HSPA1A*. Corresponding phase contrast images are shown below the fluorescence images. Scale bars represent 150 µm. **B.** Western blot on cell lysates from A detecting HSP70 and VP1 viral protein levels. Ponceau S is shown below. (*SCR*: *SCRAMBLE*; *HSP*: *HSPA1A*) **C, D.** Densitometric quantification of western blots in B. **E.** Plaque assay quantification of infectious virus in media from si*SCRAMBLE* and si*HSPA1A* transfected cells. (*< 0.05, ** *p< 0.01*; *** p<0.001; **** p<0.0001; Student’s t-test, *n* = 3).

**FIG 6.**
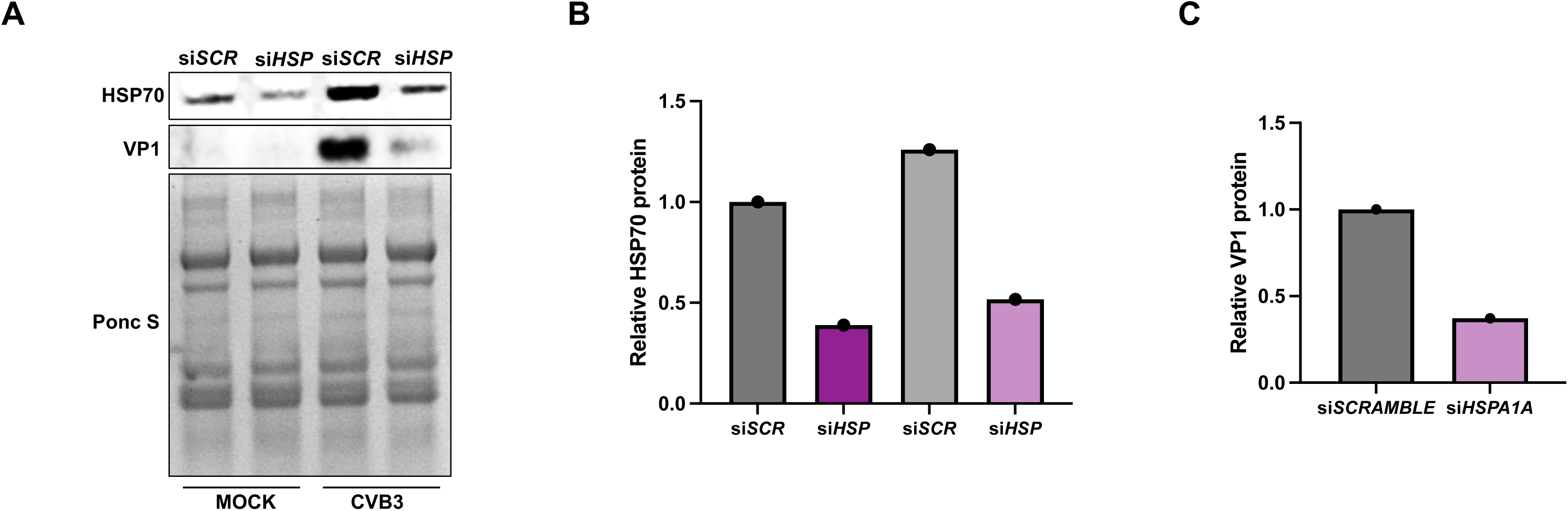
Silencing HSPA1A inhibits CVB3 release in EVs. HeLa cells were transfected with siRNA targeting HSP70 (si*HSPA1A*) or scrambled RNA (si*SCRAMBLE*) for 48 h and subsequently infected with EGFP-CVB3 at MOI 0.001 for 24 h. **A.** Western blot on isolated EVs showing HSP70 and VP1 viral capsid protein. Ponceau S is shown below. (*SCR*: *SCRAMBLE*; *HSP*: *HSPA1A*) **B, C.** Densitometric quantification of western blots in A.

### Capsazepine and SB-366791 inhibit CVB3 infection

Since capsaicin-mediated TRPV1 activation amplified infection, we investigated whether treating cells with TRPV1 inhibitors would alter CVB3 infection *in vitro*. Among the TRPV1 inhibitors, capsazepine has been one of the first discovered antagonists that were used to study TRP channels as therapeutic targets, such as in reducing chronic pain without any adverse effects [41–43]. For our studies, HeLa cells were treated with 40 µM capsazepine for 4 hours prior to CVB3 infection at an MOI of 0.001 for 48 hours. Our results indicated that suppressing TRPV1 with capsazepine significantly reduced VP1 levels (*p* = 0.0123) as well as release of virus into the media (*p* = 0.0153) (Fig. 7 A-D). However, this thiourea derivate lacks sensitivity and selectivity as it can non-specifically bind to other voltage-gated ion channels and nicotinic acetylcholine receptors. SB-366791, a cinnamide analogue, is a TRPV1 inhibitor known for its high selectivity [44]. Our previous work demonstrated the antiviral effects of SB-366791 on CVB3-infected HeLa cells [22]. To further elucidate the role of TRPV1 in CVB3 infection, HeLa cells were pre-treated with SB-366791 before undergoing capsaicin treatment and subsequent infection with CVB3 for 48 hours at an MOI of 0.001. This combination of SB-366791 and capsaicin significantly decreased viral infection, as demonstrated by reduced viral protein levels in cell lysates (*p* = 0.0016) and lower viral titers in the extracellular media (*p* = 0.0009) (Supplemental Fig. S3 A-D). These findings highlight the involvement of TRPV1 in the capsaicin-mediated exacerbation of CVB3 infection.

**FIG 7.**
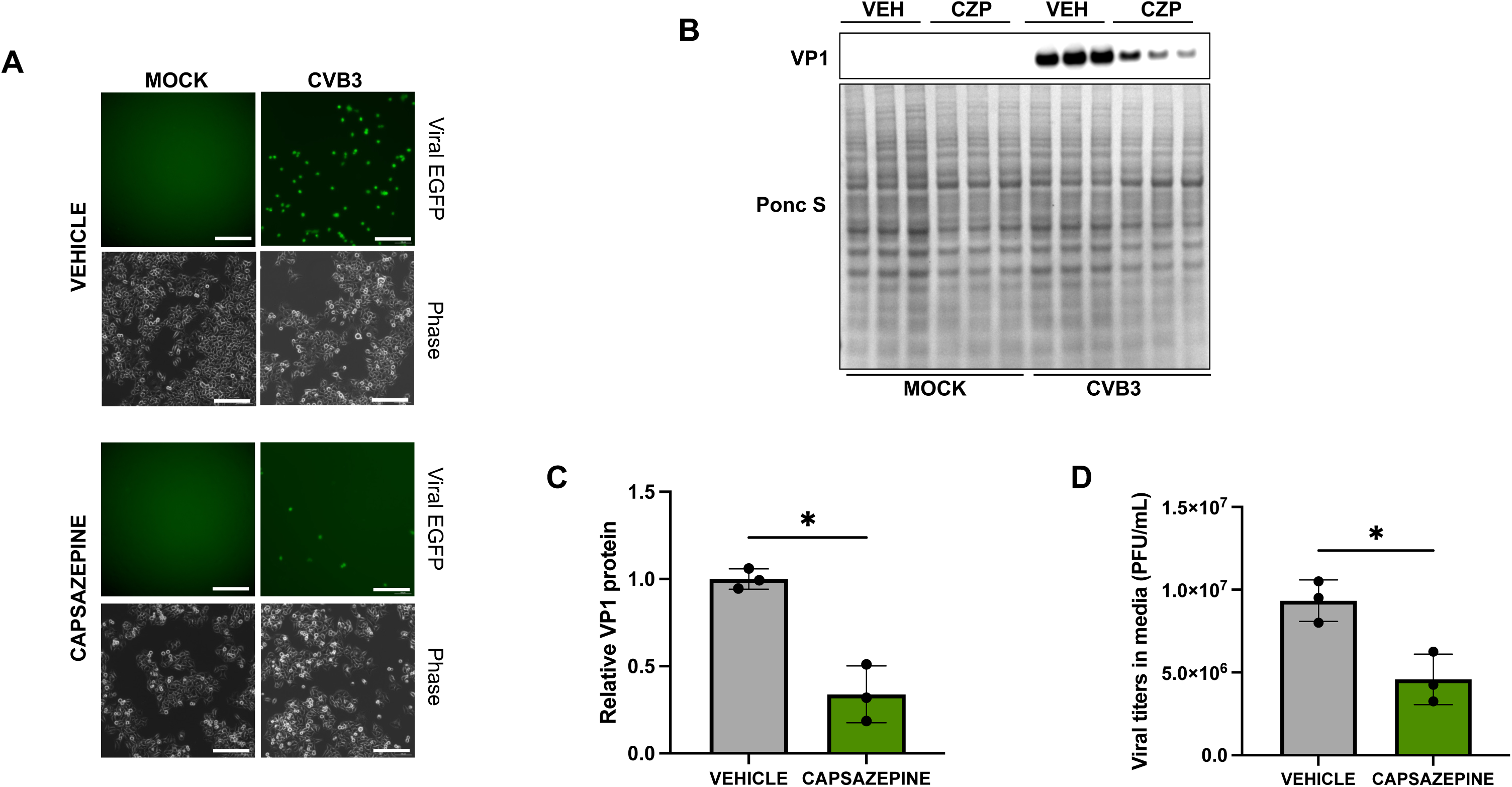
Capsazepine inhibits CVB3 infection. HeLa cells were treated with 40 µM capsazepine for 4 hours and infected with EGFP-CVB3 at multiplicity of infection (MOI) of 0.001 for 48 hours. **A.** Fluorescence microscopy compares viral EGFP expression between vehicle (DMSO-treated) and capsazepine-treated cells. Phase contrast images are shown below respective fluorescence image fields. Scale bars represent 100 µm. **B.** Western blot on cell lysates from A showing VP1 viral capsid protein. Ponceau S is shown below. (VEH: Vehicle; CZP: Capsazepine) **(C)** Densitometric quantification of western blot in B **(D)** Plaque assay quantification of infectious virus in media from cells in A. (*< 0.05, Student’s t-test, n = 3).

### SB-366791 significantly reduces CVB3 infection *in vivo*

Given that capsaicin-mediated TRPV1 activation increased infection, we sought to determine whether blocking TRPV1 would affect CVB3 infection *in vivo*. Here, we investigated the effects of SB-366791 in a mouse model of CVB3 pancreatitis. Male C57BL/6 mice were administered 1 mg/kg SB-366791 or an equivalent volume of vehicle intraperitoneally for three consecutive days. One day after treatment, mice were infected intraperitoneally with EGFP-CVB3 at a dose of 10^7^ PFU. Two days post-infection, the pancreata were harvested to perform plaque assays and histology. Throughout the treatment and infection period, the mice showed no adverse symptoms, such as reduced feeding or grooming. Plaque assay of pancreatic tissues revealed that SB-366791 significantly reduced viral titers in the pancreas (*p* = 0.0156) (Fig. 8 A). Hematoxylin and eosin staining showed severe tissue damage in vehicle-treated infected mice characterized by extensive immune cell infiltration, pancreatic edema, and necrosis. In contrast, while SB-366791-treated mice also exhibited pancreatic damage, the severity was notably reduced compared to the vehicle controls (Fig. 8 B). Furthermore, SB-366791-treated pancreatic tissues showed significant reductions in the number of necrotic cells (*p* = 0.0002), stromal cells (*p* = 0.0036) and pancreatic stellate cells (PSCs) (*p* = 0.0010), which are the primary effector cells in the process of fibrosis, a key pathological characteristic in pancreatic diseases such as chronic pancreatitis and pancreatic cancer (Fig. 8 C) [45, 46]. These findings support our *in vitro* results and underscore the role of TRPV1 in modulating CVB3 infection in host cells.

**FIG 8.**
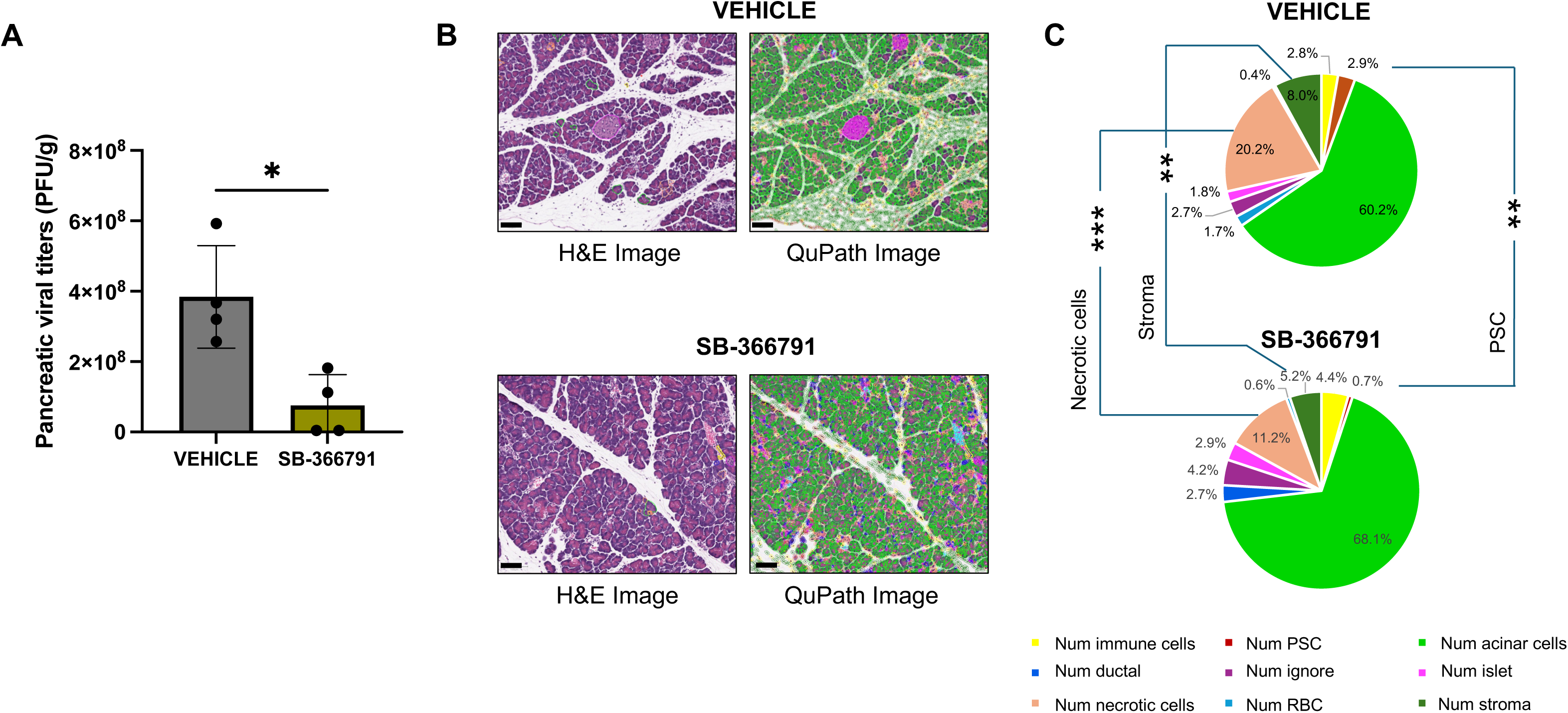
SB-366791 reduces pancreatic viral titers and tissue destruction in CVB3-infected mice. 10-week-old C57BL/6 male mice were treated with 1mg/kg menthol or equivalent volume vehicle each day for 3 days via intraperitoneal administration. On the second day of treatment, mice were infected IP with EGFP-CVB3 at a concentration of 10^7^ PFU/mouse. A total of 2 days post-infection, mice were sacrificed, and pancreata were collected. **(A)** Pancreatic viral titers as measured by plaque assays on pancreatic homogenates. (* p < 0.05; Student’s t-test, n = 4). **(B)** Representative microscopic images of the pancreas stained with hematoxylin and eosin (H&E, left panels; scale bar: 100 µm) or QuPath images (right panels; scale bar: 100 µm) for automatic quantification of cell types and stroma of same images shown in the left panels. **B)** Percentage of each cell type feature detected in vehicle (upper chart) and SB-366791 (lower chart) treated infected mice. Data are presented as pie charts (mean %). ** p≤0.005, *** p≤0.0005, all others were >0.05 and not significant.

## DISCUSSION

In the past few years, most studies have highlighted the role of TRPV1 in regulating irritant-induced airway responses and the dissemination of airborne viral particles, particularly for respiratory viruses such as rhinovirus, respiratory syncytial virus, and measles virus [47]. However, few studies have investigated the link between TRP channels and CVB3 infection. Our recent study demonstrated that the cold and menthol-sensitive transient receptor potential channel TRPM8 plays a crucial role in inhibiting mitochondrial fission (a prerequisite for mitophagy) during CVB3 infection [22]. In contrast, the role of heat-sensing ion channel TRPV1 in CVB3 infection remains understudied.

In the present study, we found that activating TRPV1 with the commonly consumed spice compound capsaicin significantly enhances CVB3 infection in HeLa cells. In addition, it markedly increases the release of CVB3-laden EVs, which have the potential to intrinsically enhance subsequent rounds of infection in neighboring cells. For many years, the *Picornaviridae* family of naked viruses were thought to escape from the host cell exclusively via cytolysis. [48–51]. This mode of dissemination releases free naked virions, making them susceptible to host neutralizing antibodies. However, since 2013, several studies have shown that certain nonenveloped viruses can utilize EVs for dissemination, thus evading host neutralizing antibodies [21, 23][52–57]. Our previous work, along with other studies have shown that CVB3 can escape host cells by packing multiple virions in phosphatidylserine-enriched membrane-enclosed EVs, thus facilitating the transfer of a high number of viral quasi-species, which can eventually infect and propagate viral progeny to neighboring cells more efficiently than free viruses [15, 30].

Upon investigating the composition of EVs released after capsaicin treatment, we observed a marked increase in the viral capsid protein VP1, as well as elevated expression of the mitochondrial protein TOM70 and the mitochondrial fission protein phospho-DRP1 (Ser 616). These findings suggest that capsaicin-mediated TRPV1 activation may promote proviral mitophagy to aid in the release of viral particles through EVs that contain mitochondrial components. In addition to these mitochondrial markers, we also observed a significant increase in the expression of HSP70 in the released EVs. The heat shock response is a highly conserved cellular response that is initiated by an increase in the fluidity of specific membrane domains, thus triggering the activation of heat-shock gene expression to safeguard and restore labile proteins and membranes. In a previous study, TRPV1 was demonstrated to act as a key regulator of the cellular thermosensory pathway, leading to membrane-dependent activation of HSPs [28]. Our findings reveal that capsaicin enhances HSP70 expression levels that exclusively increase in EVs, rather than in the cells. This suggests a previously undescribed mechanism in which HSP70 may play a pivotal role in promoting the dissemination of infectious EVs among host cells.

Earlier studies have already highlighted the proviral nature of HSP70 in the context of CVB3. This is largely due to the presence of an IRES region within the 5’UTR of the *HSPA1A* mRNA, similar to CVB3 genomic RNA, which allows it to be translated during CVB3 infection even when the cap-dependent translation is impacted [27, 58]. In our study, silencing HSP70 in HeLa cells resulted in a marked reduction of CVB3 infection, particularly in EVs, confirming the chaperone’s proviral role and supporting its importance in facilitating CVB3 spread via EVs. However, the precise role of HSP70 in the induction of mitophagy-derived EVs requires further investigation. Previous studies have linked HSP70 to both cellular autophagy and PINK1-mediated mitophagy, and HSP70 inhibition has been demonstrated to influence mitochondrial dynamics in the absence of mitophagy [40, 59]. Thus, silencing HSP70 in our study may have impeded the mitophagy process, which in turn reduced the release of infectious EVs. These findings may open new avenues for the development of antiviral therapies aimed at disrupting CVB3 transmission via EVs. Furthermore, it remains unclear exactly how the virus interacts with or influences players in this TRPV1/HSP70 pathway of EV biogenesis.

We observed that inhibiting TRPV1 with specific TRPV1 antagonist SB-366791 reduces infection *in vivo*. Earlier studies that have implicated TRPV1 in the context of pancreatitis have shown that TRPV1 levels are elevated in non-infectious animal models of pancreatitis [60, 61]. This alludes to the fact that activation of pancreatic TRPV1-positive nociceptive neurons contributes to neurogenic pancreatic inflammation. In our study, we corroborated this by finding that treating mice with TRPV1 inhibitor SB-366791 markedly attenuates CVB3 titers and tissue destruction in the pancreas.

In summary, our findings reveal a novel mechanism by which capsaicin enhances the spread of CVB3. Capsaicin-mediated TRPV1 activation induces mitochondrial fission, which leads to mitophagy. CVB3 hijacks these mitophagosomes to form mitophagosome-derived EVs, within which the virus multiplies. These EVs then facilitate viral dissemination, serving as a vehicle for CVB3 release while also aiding in immune evasion. Notably, we also observed an increase in heat shock protein HSP70 levels in capsaicin-treated EVs, suggesting a role for HSP70 in facilitating the release of infectious viral EVs (Fig. 9). Inhibiting HSP70 espression significantly impaired CVB3 release via EVs, indicating that HSP70 is crucial for viral egress. Previous studies have shown a prominent role of HSP70 in mitophagy, particularly in HEK293 cells, but whether HSP70 interacts with mitochondrial proteins or primes cells for mitophagy in the context of CVB3 infection warrants further investigation. These findings provide valuable new insights into how host cellular pathways can be co-opted by the virus to promote dissemination and immune evasion.

**FIG 9.** Capsaicin has a significant impact on CVB3 infection dynamics. **A.** Capsaicin-mediated TRPV1 activation induces mitochondrial fission, which leads to mitophagy. CVB3 hijacks these mitophagosomes to form mitophagosome-derived EVs, within which the virus multiplies. These EVs then facilitate viral dissemination, serving as a vehicle for CVB3 release. **B.** In addition, these capsaicin-treated EVs also showed higher levels of HSP 70. Earlier literature suggests that TRPV1 plays a crucial role in controlling the cellular heat shock response via activation of Heat shock Factor-1 (HSF-1) [28]. HSF-1 then trimerizes and translocate to the nucleus where it binds to *HSPA1A* promotor, resulting in translation of the protein HSP70 [65]. In our study, we observed that inhibiting HSP70 significantly impaired CVB3 release via EVs, indicating that HSP70 is crucial for viral egress. **C.** Previous studies have shown a prominent role of HSP70 in mitophagy, particularly in HEK293 cells, but whether HSP70 interacts with mitochondrial proteins or primes cells for mitophagy in the context of CVB3 infection remains to be fully explored and needs further investigation.

The fact that capsaicin is a widely consumed dietary compound raises important questions about how common dietary factors could modulate viral pathogenesis and contribute to variability in disease outcomes. Our data suggest that TRPV1 activation, through dietary exposure or pharmacological means, may significantly impact CVB3 infection dynamics. Furthermore, the identification of TRPV1 as a modulator of CVB3 spread offers promising therapeutic potential for mitigating severe disease manifestations, particularly through targeted TRPV1 inhibition. As we move forward, it will be crucial to explore how dietary interventions or TRPV1-targeted therapies could be leveraged to reduce viral burden and improve clinical outcomes in CVB3 and potentially other viral infections associated with EV-mediated transmission.

## MATERIALS AND METHODS

### Cell culture

HeLa RW cells derived from human cervical cancer cells (originally obtained from Rainer Wessly, University of San Diego, La Jolla, CA, USA) were maintained in Dulbecco’s modified Eagle’s medium (Sigma-Aldrich, D6429) supplemented with 10% FBS (Avantor, 89510-166) and antibiotic/antimycotic solution (Sigma-Aldrich, A5955). All chemicals used were of the highest grade available.

### Treatments

HeLa cells were treated with TPRV1 agonist capsaicin (Sigma-Aldrich, M2028), TRPV1 antagonist capsazepine (Sigma-Aldrich, C191) or TRPV1 antagonist SB-366791(Sigma-Aldrich, S0441). All three chemicals were dissolved in DMSO. Cells were treated with 10 μM capsaicin for 15 mins, 40 μM capsazepine for 4 hours or 10 μM SB-366791 for 24 hours prior to CVB3 infection.

### Generation of CVB3 stocks

CVB3 stocks were generated from the pMSK1 plasmid that has been generously gifted by Dr. Ralph Feuer at the San Diego State University, CA, USA. Recombinant CVB3 expressing enhanced green fluorescent protein (EGFP-CVB3) were prepared following a protocol described previously [62]. In order to generate the pMSK1 plasmid, HeLa RW cells were transfected with plasmid pH3, that encodes a myocarditic strain of CVB3 [51, 63]. A unique SfiI restriction site was then added to the backbone of this viral clone. The resulting plasmid, now known as pMSK1, can facilitate the insertion of foreign DNA fragments of interest into CVB3 genome, resulting in subsequent isolation of recombinant coxsackievirus after its transfection into HeLa cells [62]. The DNA sequence encoding enhanced green fluorescent protein (EGFP) was amplified from plasmid pEGFP and cloned into pMSK1 to generate recombinant EGFP-CVB plasmid. HeLa RW cells were subsequently transfected with the EGFP-CVB construct using Lipofectamine 2000 (Thermo Fisher). After 48 hours, cells were harvested, and freeze fractured thrice to release virions. Free-thawed cells were then centrifuged at 600 x *g* for 10 mins. The supernatant was collected and considered as ‘passage 1’ of viral stock, which were subsequently infected in fresh HeLa cells to generate ‘passage 2’ viral stocks. Concentration of the newly prepared viral stocks were then determined by plaque assay.

### CVB3 infection of host cells

HeLa cells were infected with EGFP-CVB3 at a multiplicity of infection (MOI) of 0.001. Cells were inoculated with frozen viral stocks whose viral load has been calculated by plaque assay. Equivalent amounts of DMEM growth media were added to the corresponding controls, commonly referred to as mock infected cells. After 24 or 48 hours of infection, mock and CVB3-infected cells were imaged using a Nikon Ti2 inverted epifluorescence microscope that is equipped with a Qi2 camera (Nikon, Tokyo, Japan) and an NiS Elements software version AR.5.30.05.

### Cell lysis and western blot

Following treatment and infection, all cells were lysed using chilled RIPA buffer containing 50 mM Tris-HCl (Sigma-Aldrich, 10812846001), 1% NP-40 (Sigma-Aldrich, 74385), 0.5% sodium deoxycholate (Sigma-Aldrich, D6750), 0.1% sodium dodecyl sulfate (Sigma-Aldrich, L3771), 150 mM sodium chloride (Sigma-Aldrich, 13423), 2 mM EDTA (ethylenediaminetetraacetic acid) (Sigma-Aldrich, E9884), protease inhibitor cocktail (Thermo Fisher Scientific, A32965) and PhosSTOP phosphatase inhibitors (Sigma-Aldrich, 4906845001). Cells were then mechanically disrupted, and cell slurries were centrifuged at 15000 x *g* for 10 min at 4°C and supernatants were collected to measure the protein concentrations using bicinchoninic acid solution (Thermo Fisher Scientific, 23228). Laemmli sample buffer containing 1% bromophenol blue (Allied Chemical, 0332), 1.5 M Tris-Cl pH 6.8, glycerol (Sigma-Aldrich, G5516), and β-mercaptoethanol (Thermo Fisher Scientific, BP176-100) was then added to equal amounts of protein and then loaded onto 4–12% Bis-Tris protein gels. Proteins were then transferred to nitrocellulose membranes, and membranes were stained with Ponceau S solution (Sigma-Aldrich, SLCQ5486) prior to blocking with 3% bovine serum albumin (BSA) (Sigma-Aldrich, A9647) in Tris-buffered saline with Tween 20 (TBS-T) (Sigma-Aldrich, P1379). After blocking at room temperature for one hour, the membranes were incubated with primary antibody solutions diluted in 3% Bovine Serum Albumin and TBS-T. Primary antibodies used in this study were as follows: VP1 (Mediagnost, Cox mAB 31A2, 1:2000), TOM70 (Proteintech, 14528-1-AP, 1:1000), phospho-DRP1(Ser 616) (Cell Signaling Technology, 3455, 1:1000), HSPA1A (Thermo Fisher Scientific, PA5-34772, 1:5000). Following overnight incubation, membranes were washed in TBS-T and incubated in anti-mouse secondary antibody (Sigma-Aldrich, 12-349, 1:3000) and anti-rabbit secondary antibody (Sigma-Aldrich, F9887, 1:3000). Following incubation for 1 hour, membranes were washed in TBS-T and imaged with SuperSignal West Dura Extended Duration chemiluminescent substrate (Thermo Fisher Scientific, 34075) via iBright FL1500 Imaging System (Thermo Fisher Scientific) and Azure 600 Western Blot Imaging System (Azure Biosystems). Densitometry was performed using ImageJ software (The National Institutes of Health (NIH), https://imagej.nih.gov/ij/) with background subtraction applied to all quantifications.

### Plaque assay

Following infection with CVB3, the viral titers in extracellular media were calculated using plaque assay. For this, HeLa cells were initially grown to confluency in 6 well plates. After 48 hours, media was aspirated and 400 µL of serially diluted extracellular media (from infected samples) were added to the cell monolayer. Cells were then incubated for 1 hour with occasional shaking at an interval of 15 mins. Following this, infected cells were overlaid with a 50:50 mixture of 1.2% molten agar combined with 2× DMEM. The plates were incubated at 37 °C for 48 h and agar plugs were subsequently fixed with plaque fixing solution containing 25% acetic acid and 75% methanol for 30 mins. After removing the plugs, the fixed cells were stained with 2.34% crystal violet solution for one hour following which, the plaques were counted.

### Flow Cytometry

HeLa cells were treated with 10 mM capsaicin for 15 mins prior to CVB3 infection for 48 hours at MOI of 0.001. The supernatants from infected cells were individually collected in 15 mL conical tubes. The cells were then washed with phosphate buffer saline (PBS) which was also collected in corresponding tubes to ensure collection of already detached cells. Adherent cells were then detached with Trypsin-EDTA (Thermo Fisher Scientific, 25200056) and subsequently collected in respective tubes. Following this, 10 mL growth medium was added to each tube and centrifuged at 500 x *g* for 10 minutes. The supernatant was discarded, and cells were resuspended and fixed in 5 mL 4% formaldehyde. After fixation for 15 mins, cells were centrifuged at 500 x *g* for 10 minutes. Supernatant was discarded and cells were washed by resuspending in 10 mL PBS and centrifuging at 500 x *g* for 10 minutes. Supernatant was discarded and cells were resuspended in 500 µL fresh PBS. Cells were then filtered in filter tubes to ensure single cell suspension and analyzed via Attune NxT flow cytometer (Thermo Fisher Scientific, Waltham, MA, USA).

### Isolation of extracellular vesicles

EVs were isolated from HeLa cells using the Exoquick-TC polymer-based exosome precipitation kit (Cayman, 702420). Cells were seeded in 10 cm dish, treated with capsaicin, and then infected with eGFP-CVB3 for 48 hours. One day post infection, the supernatants were collected and centrifuged at 3000 x *g* for 15 mins to eliminate residual cell debris. Supernatants were then transferred to new conical tubes, with one-fifth volume of Exoquick-TC added to each one. The tubes were incubated overnight at 4°C and EVs were pelleted by centrifugation at 1500 x *g* for 30 minutes. Purified EV pellets were washed twice with 2 mL PBS and finally harvested in chilled RIPA buffer containing protease and phosphatase inhibitors. Concentration of the EVs were measured by BCA assay and samples were subsequently prepared for western blot analysis.

### siRNA transfection

HeLa cells were seeded in 6 well plates and transfected with *HSPA1A* siRNA (Human; Santa Cruz Biotechnology, sc-29352) using Effectene Transfection Reagent (Qiagen, 301425). The transfection reagents were added following manufacturer’s guidelines. After 24 hours, the media was replenished with fresh DMEM growth media and incubated further for 24 hours before infection with CVB3 (MOI 0.001). EVs were then isolated from these cells and western blots and plaque assays were subsequently performed.

### Mouse treatments

#### Animal ethics

All experiments involving mice were performed following the National Institutes of Health guidelines and were approved by the Institutional Animal Care & Use Committee (IACUC) of The University of Alabama. The animals were anesthetized using isoflurane and were sacrificed by performing cervical dislocation.

#### Treatment

Prior to treatment, SB-366791 was dissolved in DMSO at a concentration of 10mM. Following this, the TRPV1 inhibitor was first injected intraperitoneally in 10-week old male C57BL/6 mice at 1 mg/kg/day for 2 days. At the second day of treatment, mice were infected with EGFP-CVB3 (10^7^ PFU/mice) via intraperitoneal injection. They were again treated with SB-366791 for two consecutive days post infection and were sacrificed on second day post infection. The pancreata was harvested and pancreatic tissues were either flash frozen for plaque assays or fixed with 4% formaldehyde for histology. In order to perform plaque assay, the frozen pancreatic tissue was weighed and homogenized in DMEM using a TissueLyzer LT instrument (Qiagen, Hilden, Germany). Homogenates were then centrifuged at 1000 x *g* for 10 min at 4 ^◦^C and the supernatants were collected for plaque assay.

### Histology

In order to perform histology, the fixed pancreatic tissues were paraffin embedded and subsequently sectioned into 5 µm-thick sections. These tissue sections were then deparaffinized with xylene, gradually rehydrated in ethanol, and finally stained with Gill 2 Hematoxylin and Eosin-Y following manufacturer’s protocols. The sections were then dehydrated in increasing concentrations of ethanol, washed in xylene, dried and coverslipped with Cytoseal Mounting Medium. For the quantification of distinct features including intact acinar, necrotic acinar, ductal, islet, immune and pancreatic stellate cells (PSC) and stroma, H&E slides were scanned using the Aperio AT2 microscope slide scanner (Leica Biosystems, Deer Park, IL) and imported into QuPath (open source, version 0.2.1) [64]. Each pancreas segment was captured as a single object with all cells detected. Object classification was performed with training for ∼7-8 captures per class in each slide to classify cell types and stroma, extended to all infected mice treated with vehicle (11 from 4 mice) and SB-366791 (13 from 4 mice). Ductal luminal contents were classified as “ignore”. Overall, >1 x 10^6^ cells were analyzed per group.

### Statistics

All statistical analyses were performed using GraphPad Prism version 10.2.1. For pair-wise comparison Student’s t-tests were performed and for multiple comparisons, one-way ANOVA was performed. A p < 0.05 was accepted as statistical significance. (* *p*< 0.05, ** *p*< 0.01; *** *p*<0.001; **** *p*<0.0001). *p* values >0.05 are not significant and are not labelled in the figures.

## ACKNOWLEDGEMENTS

JS is supported by National Institutes of Health (NIH) grant numbers R01DK125692 and R21AI145356. BJK is supported by NIH grant numbers R15NS131921 and R03AI185593. DF is supported by NIH grant numbers R01 HL164520, R21 AI145356, R21 AI154927. The authors would like to acknowledge Ralph Feuer (San Diego State University, CA, USA) for generously providing HeLa RW stocks and the pMKS1 plasmid and Skylar Caraway (University of Alabama, Tuscaloosa, AL, USA) for assisting in tissue culture maintenance and preparation.

## SUPPLEMENTARY MATERIALS

**FIG S1.** Histogram analysis for flow cytometry in Fig. 1B. Flow cytometry gates for quantification were defined using the termination of the fluorescent peak, at the point where the fluorescent signal was closest to zero, in a mock uninfected sample. This allowed for effective measurement of a shift in intensity or appearance of an infected population.

**FIG S2.** *Effect of capsaicin on intracellular expression of mitochondrial markers.* HeLa cells were treated with 10 µM Capsaicin for 15 mins and infected with EGFP-CVB3 at multiplicity of infection (MOI) of 0.001 for 48 h. **A.** Western blot on cell lysates detecting TOM70 and phospho-DRP1 (Ser 616). Ponceau S is shown below. (VEH: Vehicle; CAP: Capsaicin). **B, C.** Densitometric quantification of western blots in A. (*< 0.05, ** *p< 0.01*; Student’s t-test, *n* = 3).

**FIG S3.** *Combination of capsaicin and SB-366791 reduces CVB3 infection*. HeLa cells were first treated 10 µM SB-366791 for 24 hours. The following day, cells were treated with 10 µM capsaicin for 15 mins and infected with EGFP-CVB3 at multiplicity of infection (MOI) of 0.001 for 48 hours. **A.** Fluorescence microscopy compares viral EGFP expression between vehicle treated cells. Phase contrast images are shown below respective fluorescence image fields. Scale bars represent 100 µm. **B.** Western blot on cell lysates from A showing VP1 viral capsid protein. Ponceau S is shown below. (VEH: Vehicle; CAP: Capsaicin; SB: SB-366791; COMBO: Combination of capsaicin and SB-366791) **C.** Densitometric quantification of western blot in B. **D.** Plaque assay quantification of infectious virus in media from cells in A. (*< 0.05, ** *p< 0.01*; *** p<0.001; **** p<0.0001; Student’s t-test, *n* = 3).

